# Comparative gene editing reduces dopamine receptor levels across rodent species

**DOI:** 10.1101/2025.10.21.683401

**Authors:** Sonia C. Karkare, Dario Aspesi, Keith M. Garner, Evelien H.S. Schut, H. Elliott Albers, Frank J. Meye, Malavika Murugan, Arjen J. Boender

## Abstract

Translational challenges in neuroscience originate from species-specific differences that limit the generalizability of experimental findings. Comparative approaches can help distinguish conserved from species-specific mechanisms, but their application has been limited by the lack of molecular tools beyond traditional model organisms, complicating direct comparisons of conserved and divergent mechanisms of neural function. This gap is particularly evident for the dopaminergic system, a key regulator of motivated behaviors across species and the principal pharmacological target for current psychotherapies. Building on our recent development of comparative gene editing, we here present an adeno-associated virus-mediated CRISPR/Cas9 strategy to reduce *in vivo* dopamine receptors D1 and D2 levels across the rodent phylogeny. Using this approach, we achieved specific reduction of receptor levels in three rodent species (house mouse, prairie vole, and Syrian hamster), which we demonstrate with radioactive ligand binding assays. This toolkit expands the reach of comparative gene editing approaches, enabling functional investigation of the dopaminergic system across rodent species. Thereby, it supports comparative neuroscience by facilitating the identification of conserved versus species-specific neural mechanisms with enhanced translational potential.

## Introduction

The diversity of nervous systems across species offers a powerful framework for understanding the mechanistic underpinnings of behavior (Beach, 1950). Comparative neuroscience uses this natural variation to identify both conserved principles of neural circuit function and species-specific adaptations shaped by ecological demands (Johnson and Young, 2018; Mathuru et al., 2020; Mars et al., 2021). Progress in this field, however, has been limited because most molecular and genetic tools are optimized for a narrow set of traditional laboratory models. While pharmacological, optogenetic, and chemogenetic approaches can and have been applied across species, their capacity to interrogate GPCR function is limited by off-target effects and the inability to selectively manipulate receptor subtypes in defined neural circuits. Recent advances in genome sequencing, viral delivery, and gene editing technologies have now made it feasible to apply genetic approaches to a broader range of organisms (Bengston et al., 2018; Luo et al., 2018; Boender and Young, 2020; Moulin et al., 2021). These developments enable us to expand investigations beyond standard rodent models like house mice and Norway rats and to examine how neural circuits generate behavioral diversity across the rodent phylogeny.

In previous work, we introduced an adeno-associated (AAV)–CRISPR/Cas9 system to reduce oxytocin receptor (OXTR) levels across multiple rodent species, demonstrating the feasibility of cross-species, or comparative gene editing (Boender et al., 2023). Here, we extend this approach to the dopamine (DA) system, a core neurotransmitter system central to movement, reward, motivation, and cognition (Costa and Schoenbaum, 2022). DA receptors fall into two families: DRD1-like (DRD1, DRD5), which activate adenylate cyclase and increase cAMP, and DRD2-like (DRD2, DRD3, DRD4), which inhibit this pathway and reduce cAMP (Boyson et al., 1986; Gerfen et al., 1990; Beaulieu et al., 2015). While the DA system is broadly conserved across vertebrates (Calipari et al., 2012), DA receptor subtypes differ subtly in their regional expression within and across species, supporting both conserved and divergent behavioral mechanisms (Missale et al., 1998; Yamamoto et al., 2013; Yuan and Zhao, 2020). Moreover, genetic variation in DA receptor genes has been implicated in numerous neuropsychiatric and neurodegenerative conditions, including schizophrenia, substance use disorders, Parkinson’s disease, and Alzheimer’s disease (Rampino et al., 2018; Magistrelli et al., 2021; Speranza et al., 2025). As social disability often represents the earliest manifestation of psychiatric conditions characterized by DA dysfunction (Porcelli et al., 2019; Holt-Lunstad, 2022), we believe that comparative studies of DA signaling across species with diverse social behaviors are essential for understanding the neural basis of mental Illness (Phelps et al., 2010).

Despite the translational relevance of intra- and interspecific variation in the DA system, functional studies remain heavily focused on isogenic house mouse and Norway rat strains. Other rodents, however, exhibit DA-dependent social behaviors that are difficult to study in these standard models yet are highly relevant to the human condition. Prairie voles, for instance, have revealed how DA signaling in the nucleus accumbens () regulates pair bonding behaviors: DRD2-like activity promotes bond formation, whereas DRD1-like signaling opposes the formation of new pair bonds but promotes pair bond maintenance (Gingrich et al., 2000; Aragona et al., 2003, 2006). Interestingly, DA signaling does not seem to be crucial for same-sex, or peer bonds, suggesting that different types of social bonds are controlled by diverging underlying mechanisms (Lee and Beery, 2021).

In Syrian hamsters, DA signaling also contributes to aggression, stress responses, and mating through circuits that integrate with nonapeptide and chemosensory pathways (Gray et al., 2015; Morrison et al., 2015; Cross et al., 2025). While DA involvement in aggression and stress is conserved across rodents, hamsters provide a unique opportunity to study these processes in both sexes, as females are as aggressive as—or even more aggressive than—males (Elidio et al., 2021). In house mice and Norway rats, DA signaling has been extensively studies and contributes to a multitude of neural functions and behavior, including the processing of social information (Bariselli et al., 2018; Dai et al., 2022; Solié et al., 2022), maternal care (Champagne et al., 2004; Xie et al., 2023), stress-induced behaviors (Linders et al., 2022) and juvenile play (Manduca et al., 2016; Vanderschuren et al., 2016). Because DA signaling underlies both conserved and species-specific social behaviors of translational relevance, there is a critical need for comparative tools that reveal how this system shapes behavioral variation. Such approaches are essential for understanding how DA signaling contributes to human sociality in both health and disease.

In the present study, we designed and validated an AAV–CRISPR/Cas9 system that selectively reduces DRD1 and DRD2 levels in three rodent species: *Mus musculus* (house mouse), *Microtus ochrogaster* (prairie vole), and *Mesocricetus auratus* (Syrian hamster). By targeting conserved coding regions, we achieved reliable *in vivo* reduction of DA receptor levels across all three species, which we confirm by receptor autoradiography. This approach establishes a comparative neuroscience framework for selective DA receptor manipulations, enabling new investigations into the role of DA signaling in translationally relevant social behaviors such as pair bonding, maternal care, play, aggression, and mating.

## Results

### Design of viral strategy to reduce dopamine receptor levels in multiple rodents

We sought to engineer a viral vector-based approach for reducing dopamine receptor (DRD1 and DRD2) levels in brain tissue across diverse rodent model systems. Our strategy involved testing the effectiveness of this tool in three rodent species commonly employed in the study of social behavior: *Mus musculus* (house mouse), *Microtus ochrogaster* (prairie vole) and *Mesocricetus auratus* (Syrian hamster). Previously, we established an AAV-CRISPR/Cas9 methodology employing a guide RNA (gRNA) directed against conserved regions in rodent *Oxtr* genes, which successfully and specifically diminished functional OXTR levels in six rodent species (Boender et al., 2023). Here, we implemented a comparable strategy utilizing gRNAs directed against conserved sequences in rodent *Drd1* and *Drd2* genes. To pinpoint conserved DA receptor coding regions suitable for CRISPR/Cas9-mediated disruption, we employed the ClustalW alignment algorithm within the msa package of R/Bioconductor. For both receptors we identified conserved domains of adequate length for gRNA design (>19 nucleotides) that also harbored compatible protospacer adjacent motif (PAM) sequences (5′-NGG-3′) (Figure 1). The gRNA(ΔDRD1) target site is positioned within transmembrane domain 5, while gRNA(ΔDRD2) targets a sequence in the fourth transmembrane domain.

**Figure 1.**
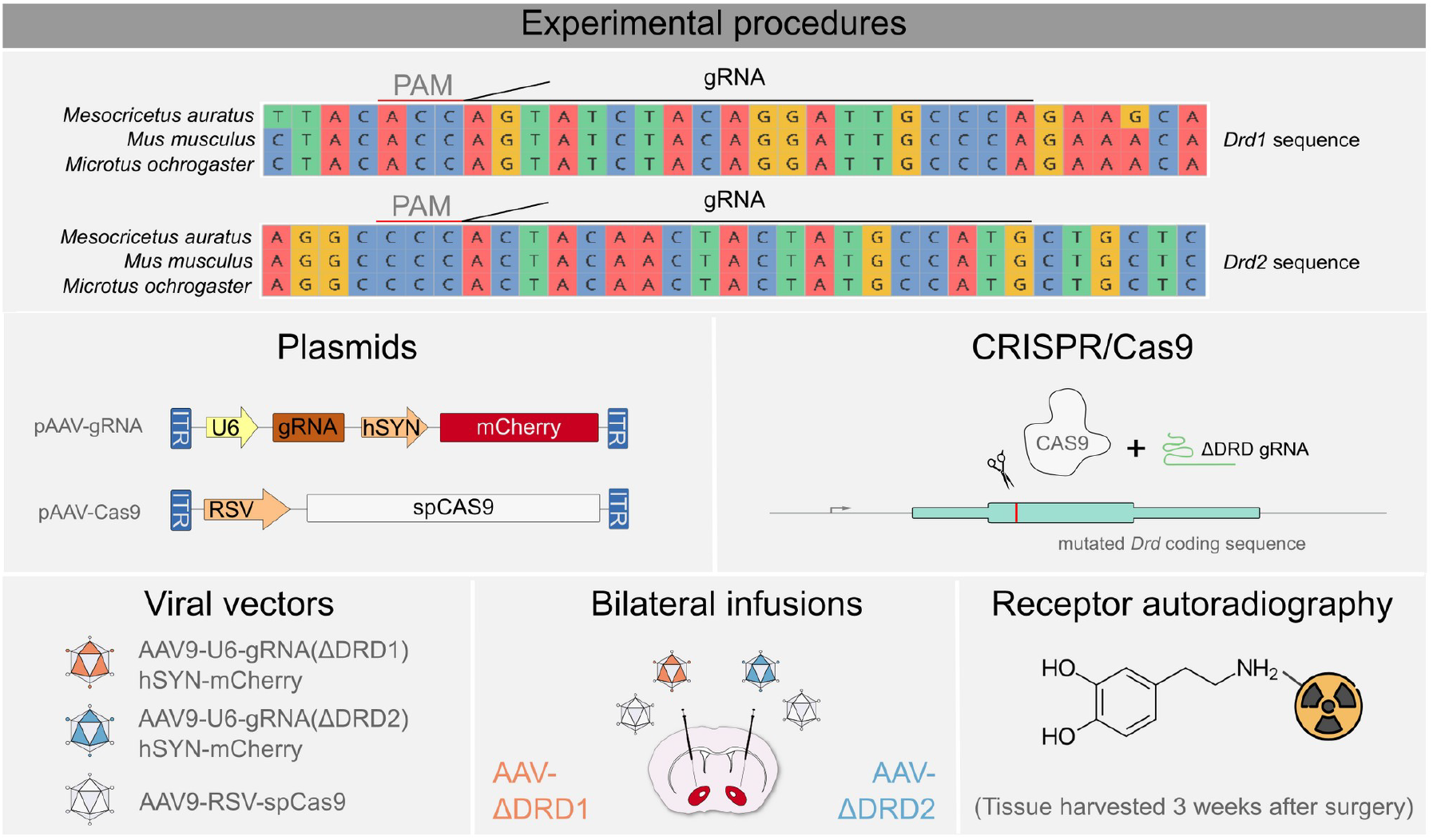
Design of AAV-CRISPR/Cas9 strategy to reduce dopamine receptor levels across rodent species. Schematics of experimental procedure that detail the conserved regions in *Drd1* and *Drd2* that are targeted by our strategy, the plasmids that were used, CRISPR/Cas9 mutagenesis, the viral vectors, the surgical approach and the validation of efficacy and selectivity of viral vectors by receptor autoradiography.

### Validation of the viral strategy in three rodent species

We generated AAV-gRNA(ΔDRD1) and AAV-gRNA(ΔDRD2) vectors to target these conserved regions and co-delivered AAV-gRNAs with an AAV-Cas9 vector. AAV-gRNA(ΔDRDs) were unilaterally injected in the NAc, whereas the opposite hemisphere received AAV-gRNA(LacZ). The latter vector targets a bacterial gene sequence absent from rodent genomes, and thus serves as a control. AAV-CRISPR/Cas9–mediated genomic editing was initially validated through a T7 endonuclease assay (Figure S1A). DNA was isolated from mCherry–positive tissue, and the AAV-CRISPR/Cas9– targeted DA receptor coding regions were polymerase chain reaction (PCR)–amplified. T7 endonuclease restriction was observed in PCR amplicons, and digested products were of the expected length in isolates from AAV-ΔDRDs–infected tissue, but not present in AAV-LacZ–infected tissue isolates (Figure S1B). This indicates that specific mutagenesis had occurred in the DA r*e*ceptor coding sequences. Subsequently, we evaluated the efficacy of AAV-gRNA(ΔDRD2), to determine whether these mutations translated into reduced DRD2 protein levels and set-up DA receptor autoradiography. Following a three-week post-surgical interval, we observed reduced DRD2 levels in AAV-gRNA(ΔDRD2) infused hemispheres in all three species, as visualized by ^3H^Raclopride autoradiography (Figure S2).

Next, we unilaterally injected AAV-gRNA(ΔDRD1) mixed with AAV-Cas9 into the ipsilateral NAc, while the contralateral hemisphere received AAV-gRNA(ΔDRD2) with AAV-Cas9 (Figure 2). This design allowed each DA receptor-targeting vector to serve as the control for the other, providing an internal measure of both specificity and efficacy within the same animal. The gRNA(LacZ), which has no known target in mammalian genomes, contains nine mismatches relative to both mouse *Drd1* and *Drd2* coding sequences. Our gRNA(ΔDRD1) and gRNA(ΔDRD2) sequences each have eight mismatches with the respective non-target DA receptor coding sequence. Previous studies indicate that even one or two mismatches, particularly in the PAM-proximal “seed” region, can markedly reduce or abolish editing efficiency (Anderson et al., 2015; Modrzejewski et al., 2020). The most likely potential off-target sequences in our study showed at least five mismatches and lacked a permissive PAM, making off-target mutagenesis extremely unlikely. Consistent with these predictions, AAV-gRNA(ΔDRD1) significantly reduced ^3H^SCH23390 levels (65.7%±4.86 in Syrian hamster, 65.5%±3.98 in house mouse, and 39.2%±5.83 in prairie vole), while AAV-gRNA(ΔDRD2) significantly diminished ^3H^Raclopride levels in the NAc of all three species (51.9%±8.03 in Syrian hamster, 45.4%±3.65 in house mouse, and 40.6%±4.12 in prairie vole). This result confirms that the AAV-CRISPR/Cas9 strategies specifically perturbed functional dopamine receptor synthesis (Figure 3). While DRD1 level was significantly reduced in prairie voles, it was significantly less reduced in comparison to the other species (*Padj*<0.001, Tukey-HSD). The observed reduction in dopamine receptor levels was not attributable to neuronal loss, as control hemispheres still exhibited radioactive binding activity, indicating specific loss of the targeted dopamine receptor.

**Figure 2.**
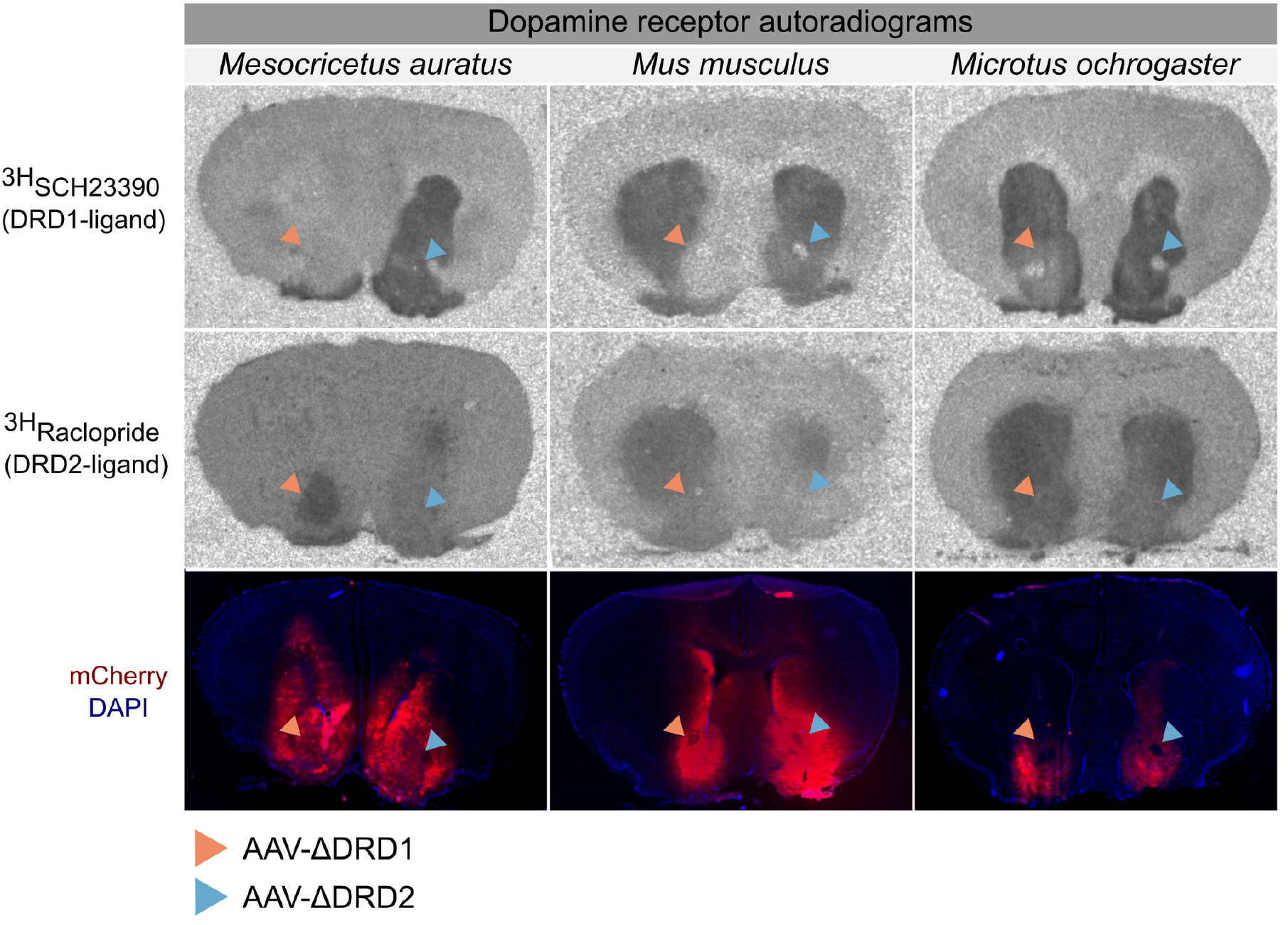
Dopamine receptor autoradiographies demonstrating the ability of the AAV-CRISPR/Cas9 strategy to reduce dopamine receptor levels across rodent species. Representative autoradiographies of ^3H^SCH23390 and ^3H^Raclopride binding, as well as images of the viral-induced mCherry expression counterstained with DAPI. All images in one column are adjacent brain sections from the same individual. Orange arrows indicate AAV-ΔDRD1 injections, while blue arrows represent AAV-ΔDRD2 infusions

**Figure 3.**
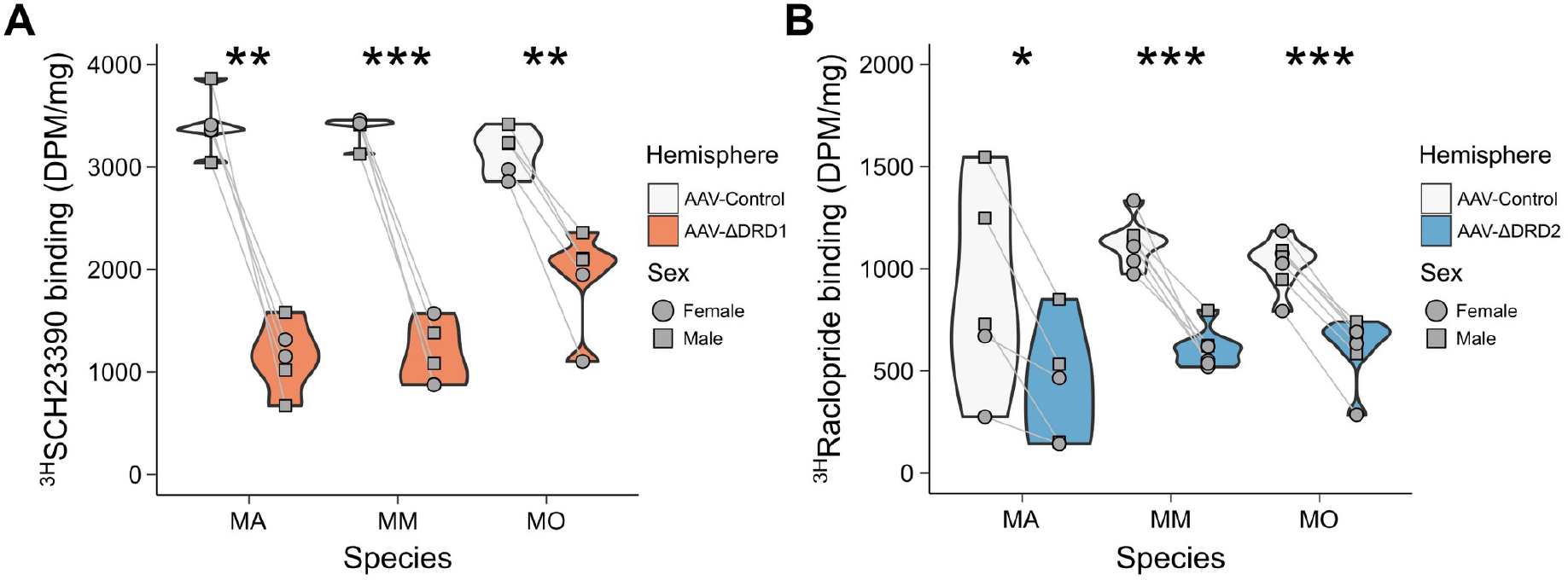
Both AAV-CRISPR/Cas9 strategies are effectively and specifically reducing dopamine receptor levels. **A)** Quantification of ^3H^SCH23390 binding in AAV-ΔDRD1 versus AAV-Control injected hemispheres. Paired *t* tests corrected for multiple comparisons: MA (Syrian hamster) - \*\**Padj*=0.003; MM (house mouse) - \*\*\**Padj*=0.001; MO (prairie vole) - \*\**Padj*=0.002. **B)** Quantification of ^3H^Raclopride binding in AAV-ΔDRD2 versus AAV-Control injected hemispheres. Paired *t* tests corrected for multiple comparisons: MA (Syrian hamster) - \*\**Padj*=0.03; MM (house mouse) - \*\*\**Padj*=0.001; MO (prairie vole) - \*\*\**Padj*=0.0001.

### Predicted effectiveness across rodent species

Finally, we searched all available rodent RefSeq RNA data for species in which gRNA(ΔDRD1) and/or gRNA(ΔDRD2) target sequences aligns perfectly to the respective DA receptor genes and are thus predicted to work. We collected available rodent RefSeq *Drd1* and *Drd2* sequences (45 species) and predict gRNA(ΔDRD1) to be functional in 26 species and gRNA(ΔDRD2) to be functional in 27 species (Figure 4). Extrapolating from these statistics, it is our estimate that these gRNAs will be functional in ∼60% of rodent species.

**Figure 4.**
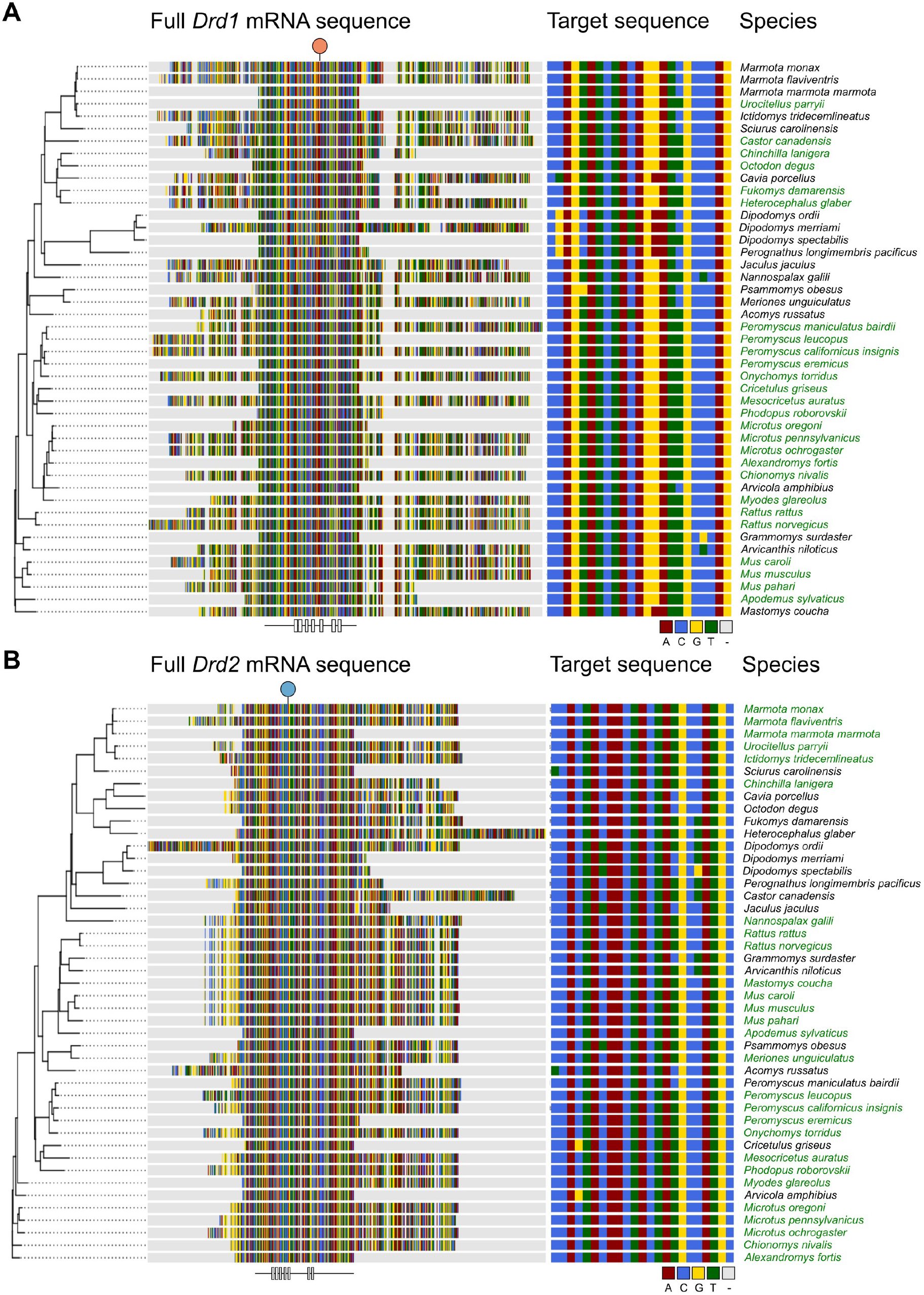
Alignment of all current rodent RefSeq DA receptor mRNA sequences and their match to the gRNA target sequences. **A)** Depicted on the left is a multiple sequence alignment of all available RefSeq rodent *Drd1* mRNA sequences (*N* = 45). The circle on top indicates the position of the gRNA target sequence, and the labels on the bottom indicate the transmembrane portions of the protein. In the middle, the gRNA target regions are shown. Scientific species names are shown on the right, green color indicates perfect alignment to the gRNA and the permissive PAM-sequence. **B)** As A but for *Drd2*.

## Discussion

We developed a flexible and effective strategy to selectively reduce DRD1 or DRD2 levels in at least three rodent species: prairie voles, Syrian hamsters, and house mice. By targeting conserved coding regions, the same AAV-mediated CRISPR/Cas9 approach produced reliable, receptor-specific reductions in NAc receptor levels without broader neuronal loss. This establishes a comparative genetic tool for dissecting DA signaling across rodent species.

Unlike systemic knockouts, which can produce developmental confounds (Fowler et al., 2002; Nakamura et al., 2014), our region-specific strategy enables targeted manipulation of mature neurons. It offers several advantages when compared to *in vivo* pharmacology. First, as pharmacological reagents often have off-target binding, the CRISPR/Cas9 strategy offers more specificity than DA receptor ligands (Butini et al., 2016; Myslivecek, 2022). Second, the strategy can be combined with other minimal promoters to target specific cell populations (Schimmer et al., 2024). Third, it is better suited for long-term behavioral experiments that require persistent manipulations (Nerio-Morales et al., 2024; Fricker et al., 2025). Finally, the approach allows for region-specific manipulation of DA-dependent signaling, and can be extended to other brain regions, such as the amygdala, cortex, hippocampus, and striatum, where DA signaling regulates reward, cognition, and addiction (Costa and Schoenbaum, 2022). Abnormalities in these circuits contribute to neuropsychiatric disorders including schizophrenia and Parkinson’s disease (Rampino et al., 2018; Magistrelli et al., 2021; Speranza et al., 2025). A versatile cross-species tool increases the likelihood of capturing both conserved and species-specific contributions of DA-receptors to behavior. For example, prairie voles provide a model for pair bonding (Walum and Young, 2018), while Syrian hamsters enable investigation of sex-dependent territorial aggression (Terranova et al., 2016), behaviors not accessible in the more standard house mouse or Norway rat models

As noted, this strategy creates opportunities to study how sex differences shape dopamine-dependent behaviors. Female hamsters are as aggressive as or more aggressive than males (Elidio et al., 2021), female prairie voles pair bond earlier than males (Schaepe and Hiura, 2025), and male and female mice differ in social and anxiety-related behaviors (Cox and Rissman, 2011; Hashikawa et al., 2018; Mariscal et al., 2023). Because these sex differences vary across species, comparative approaches are essential for identifying the molecular mechanisms that govern them. Our method enables targeted investigation of DRD1 and DRD2 receptor contributions to these behaviors in both sex- and species-dependent contexts.

A technical limitation of this approach is that it depends on sequence conservation. The three species studied here shared conserved *Drd1* and *Drd2* coding regions that permitted the use of a single gRNA design. Whether this strategy can be generalized to other species requires further testing. However, our prior work of comparative gene editing of rodent *Oxtr* genes predicts that conserved sites amenable for gene editing exist across many rodent and mammalian species (Boender et al., 2023). In a similar fashion, our BLAST search in available RefSeq rodent transcriptomes suggest that AAV-gRNA(ΔDRD1) works in at least 26 rodent species, while AAV-gRNA(ΔDRD2) should be functional in at leasr 27 rodent species. Both these vectors are predicted to work in species of the genus *Rattus, Peromyscus* and *Mus*, which are other relatively common, (non)-traditional animal species used to understand the neurobiological basis of social behavior (Trainor et al., 2011; Chen et al., 2023; Klibaite et al., 2025).

Notably, our AAV-CRISPR/Cas9 strategy did not lower DA receptor levels as effectively as it reduced OXTR levels in our previous experiments (>90% versus >40% in this study) (Boender et al., 2023), despite using the same species, viral constructs, and methodological framework. Although differences in receptor turnover rates or gRNA efficiency could play a role, the most plausible explanation lies in the pharmacological specificity of the DA receptor ligands employed here. While SCH23390 and Raclopride are potent ligands for DRD1 and DRD2 respectively, these compounds also show considerable affinity for DRD5 and DRD3 respectively (Myslivecek, 2022). Indeed, prairie voles express relevant levels of these receptors in the NAc (Loth et al., 2025), which our ligands may have bound to. As our gRNAs are not likely to target these subtypes (>5 mismatches and no PAM sequence), any residual binding after knockdown is likely to have been caused by non-specific binding of the ligands, rather than reduced efficacy of our approach.

Overall, this study establishes a cross-species framework for targeted manipulation of DRD1 and DRD2 receptors. By enabling receptor-specific knockdown in diverse model organisms, our approach expands the behavioral and evolutionary contexts in which DA signaling can be studied. Broadening comparative research may help overcome the translational gap between rodent models and human disease (Sun et al., 2022), and improve our understanding of DA contributions to motor and psychiatric disorders such as Parkinson’s disease, schizophrenia, and substance use disorders (Speranza et al., 2025). Ultimately, this flexible genetic tool provides new opportunities to link species diversity in DA signaling with behavioral diversity in both health and disease.

Our cross-species DA receptor editing approach enables the investigation of fundamental questions about the relationship between molecular conservation and functional divergence in DA circuits. While DRD1 and DRD2 expression patterns are relatively conserved across rodents (Yamamoto et al., 2013; Yuan and Zhao, 2020), their downstream signaling cascades appear to generate remarkable behavioral diversity. Comparative studies suggest that the DA machinery can be opted to produce a diverse range of behavioral outputs. For instance, DRD2 signaling in prairie vole NAc promotes pair bond formation (Aragona et al., 2006), whereas in Syrian hamsters similar circuits may regulate territorial aggression (Gray et al., 2015; Morrison et al., 2015). Investigation of such functional plasticity within conserved molecular frameworks has the potential to give new insights into how evolutionary pressures shapes circuit-level organization without altering basic neurotransmitter systems (Bendesky et al., 2017; Baier et al., 2025).

From a translational perspective, species-specific functional divergence may explain why certain therapies that target the DA system show variable efficacy across patient populations (Sun et al., 2022). Human genetic variation in DA receptor expression patterns could mirror the species-level differences we observe in rodents, potentially creating subpopulations with subtle, but distinct circuit functioning. Indeed, some therapy-resistant patient populations might reflect natural variation in DA system organization that are masked in isogenic laboratory strains.

With this work, we hope to facilitate the systematic mapping of DA circuit diversity across species, an endeavor that could inform novel approaches and therapies to modulate DA system functioning. By characterizing how variation in DA receptor signaling drives behavioral phenotypes across the rodent phylogeny, we may develop useful frameworks for precise therapeutic interventions that match individual circuit architectures. Such approaches will help advance psychiatric treatment towards personal interventions rooted in evolutionary neurobiology.

## Material and Methods

### Design and synthesis of viral vectors

*Drd1* and *Drd2* coding sequences from *Mesocricetus auratus, Microtus ochrogaster and Mus musculus* were aligned with the ClustalW algorithm in the R/Bioconductor msa package to identify conserved regions in which candidate gRNA sequences were identified using Benchling. Two gRNA sequences were selected on the basis of predicted efficiency and off-target effects. A control gRNA was used to target the bacterial LacZ gene. gRNA sequences were the following: gRNA(ΔDRD1), 5′-CTGGGCAATCCTGTAGATAC-3′; gRNA(ΔDRD2), 5′-GCATGGCATAGTAGTTGTAG-3′; and gRNA(LacZ), 5′-GTGAGCGAGTAACAACCCGT-3′. Oligos were cloned into cloned into pAAV-U6-gRNA-hSYN-mCherry (gift from Alex Hewitt, Addgene plasmid #87916) after plasmid digestion with Sap I (Hung et al., 2016). Viral particles were synthesized using pAAV-U6-gRNA-hSYN-mCherry, pAAV-RSV-spCas9 (Addgene, plasmid #85450, gifted by H. Lei) (Hung et al., 2016), pAAV9-SPAKFA (Penn Vector Core, PA, USA), and pAAV/Ad (American Type Culture Collection, VA, USA). AAV9 particles were produced in human embryonic kidney (HEK) 293T cells, purified with AVB-affinity chromatography (Wang et al., 2015), and concentrated by centrifugal filtration (Amicon Ultra-4, Fisher Scientific, NH, USA), after which viral titer was determined using quantitative PCR targeting the inverted terminal repeats. Three viruses were generated: AAV9-U6-gRNA(ΔDRD1)-hSYN-mCherry, AAV9-U6-gRNA(ΔDRD2)-hSYN-mCherry and AAV9-U6-gRNA(CTRL)-hSYN-mCherry. AAV9-RSV-spCas9 was generated by VectorBuilder (Chicago, IL, USA). AAV-gRNA and AAV-Cas9 vectors were diluted to 3.0 × 1010 and 1.5 × 1010 genomic copies/μl, respectively, and mixed in a 1:1 ratio.

### Animals

All experiments were performed following the guidelines and approved by the respective Institutional Animal Care and Use Committees (Emory University, Georgia State University and Utrecht Medical Center Utrecht). 22 animals were used in total: 6 Syrian hamsters (*Mesocricetus auratus)* at Georgia State University (3 males and 3 females), 7 house mice (*Mus musculus*) at Emory University (3 males and 4 females), 2 house mice at Utrecht Medical Center Utrecht (2 males) and 7 prairie voles (*Microtus ochrogaster*) at Emory University (3 males and 4 females). All animals were sexually naïve adults (>PND60) and housed in standard laboratory conditions with *ad libitum* water and food provided.

### T7 endonuclease I assay

mCherry-infected tissue was collected from the NAc of fresh-frozen mouse brain sections, and DNA was isolated using the Blood & Tissue DNA kit (Qiagen, Germany). Fragments surrounding the gRNA target sides were PCR-amplified using Q5-polymerase (New England Biolabs, MA, USA) and the *Drd1* primer set 5′-GAGGGACTTCTCCTTTCGCAT-3′ (forward) and 5′-GAGCATTCGACAGGGTTTCCA-3′ (reverse) and *Drd2* primer set 5′-TATGGCTTGAAGAGCCGTGC-3′ (forward) and 5′-TATGGCTTGAAGAGCCGTGC-3′ (reverse). Both sets used these cycling conditions: 98°C for 2 min, 34 cycles of 98°C for 15 s, 63°C for 15 s, and 72°C for 30 s, and 2 min of final elongation. Next, PCR fragments were used for the T7 endonuclease I assay according to the manufacturer’s instructions (Integrated DNA Technologies, IA, USA).

### Intracranial surgeries

For all species, anesthesia was induced by exposure to a 2 to 4% isoflurane/oxygen mix and maintained at 1-3%. Three daily doses of meloxicam or carprofen (2 to 5 mg/kg, depending on species) were administered after surgery. Using a stereotaxic apparatus, most animals (n=15), were unilaterally injected with a 1:1 mix of AAV9-RSV-Cas9 and AAV9-gRNA(ΔDRD1), while the contralateral side received a 1:1 mix of AAV9-RSV-Cas9 and AAV9-gRNA(ΔDRD2). The remaining animals received unilateral injection of a 1:1 mix of AAV9-RSV-Cas9 and AAV9-gRNA(ΔDRD2), while the contralateral side was injected with a 1:1 mix of AAV9-RSV-Cas9 and AAV9-gRNA(LacZ). Stereotaxic coordinates and injected volumes are summarized in Table 1.

**Table 1.**
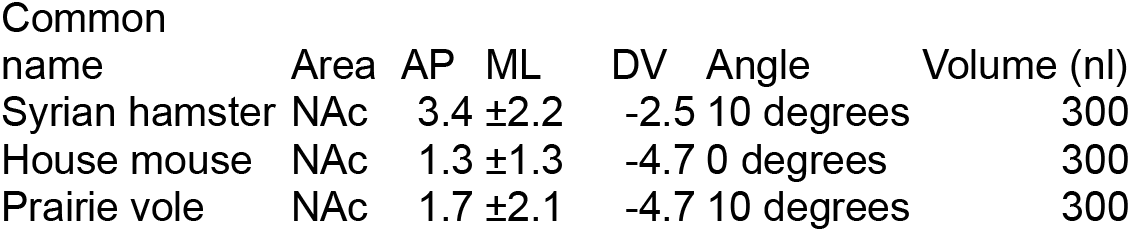
Stereotaxic coordinates used for viral injections.

### DRD1 and DRD2 autoradiography

Fresh-frozen brains were sectioned on a cryostat (Epredia Cryostar NX-70, Thermo Fisher Scientific, MA, USA) at 20 μm, mounted on Superfrost Plus slides (Fisher Scientific), and stored at −80°C until use. Autoradiography was performed as previously described (Defagot and Antonelli, 1997; Lee and Beery, 2021). Briefly, slides were thawed and pre-incubated in incubation buffer (25 mM Tris-HCL, 100 mM NaCl and 1 mM MgCl2 in PBS, pH 7.5) at RT for 20 minutes. Next, sections were incubated in buffer supplemented with either 4.5 nM of DRD1-ligand ^3H^SCH23390 (75 Ci/mmol, = #NET930250UC, Revvity, MA, USA) and 1 µM of ketanserin (to prevent binding to 5-HT2 receptors) or with 4.5 nM of the DRD2-ligand ^3H^Raclopride (75 Ci/mmol, #NET975250UC, Revvity, MA, USA) at room temperature (RT) for 2 hours. Unbound ligand was removed by incubation in ice-cold buffer (4°C) for 30 minutes and three dips in MQ. Finally, slides were placed in a cassette with phosphor imaging plates (Fujifilm, Japan) for 5 (DRD1/SCH23390) or 12 (DRD2/Raclopride) days. After exposure, plates were imaged with a BAS-5000 phospor imager (Fujifilm). Mean gray values of viral-targeted regions, corrected for background, were determined in ImageJ. Radioactivity (disintegrations per minute) was calculated using an C14-standard and taken as a proxy for DRD1 or DRD2 density. Reductions in DRD1 or DRD2 densities were determined by comparing protein density levels in the hemisphere infused with AAV-gRNA(ΔDRD) to the contralateral hemisphere that received either AAV-gRNA(LacZ) or the other AAV-gRNA(ΔDRD), collectively classified as AAV-Control.

### Statistical analyses

All statistical analyses were performed in RStudio, using paired *t* tests (Figure 2) with α = 0.05.

### BLAST

We used the blastn algorithm in the NCBI BLAST+ software suite to search for perfect alignment of the target RNA sequences plus the permissive PAM sequence (5′-NGG-3′) to rodent (taxid 9989) RefSeq mRNA to identify species in which our tools are predicted to reduce DRD1 and/or DRD2 levels.

## Author contributions

Conceptualization: A.J.B and F.J.M. Design of viral strategy: A.J.B. Synthesis of viral particles: A.J.B. and K.M.G. Injection of viral particles: A.J.B., D.A., S.C.K. Contribution of animals: F.J.M, H.E.A. and M.M. Analyses and statistics: A.J.B. Supervision: A.J.B., F.J.M., H.E.A, and M.M. Writing—draft: A.J.B. D.A., and S.C.K. Writing—review and editing: all authors. Order of first authors was determined by a coin flip.

## Acknowledgments

Our special thoughts go out to Dr. Larry Young, mentor to Dr. Arjen Boender, who was very supportive of this work and granted us the use of his resources shortly before his passing. We will always remember him for his collaborative spirit and his love for comparative neuroscience.

## Data and materials availability

The necessary AAV-gRNA and AAV-Cas9 plasmids to produce these viral vectors will be made available through Addgene upon publication. Datasets and code will be uploaded to the Dryad repository upon publication.

