## Supplementary material for "Comparative gene editing reduces dopamine receptor levels across rodent species": Figures S1 and S2

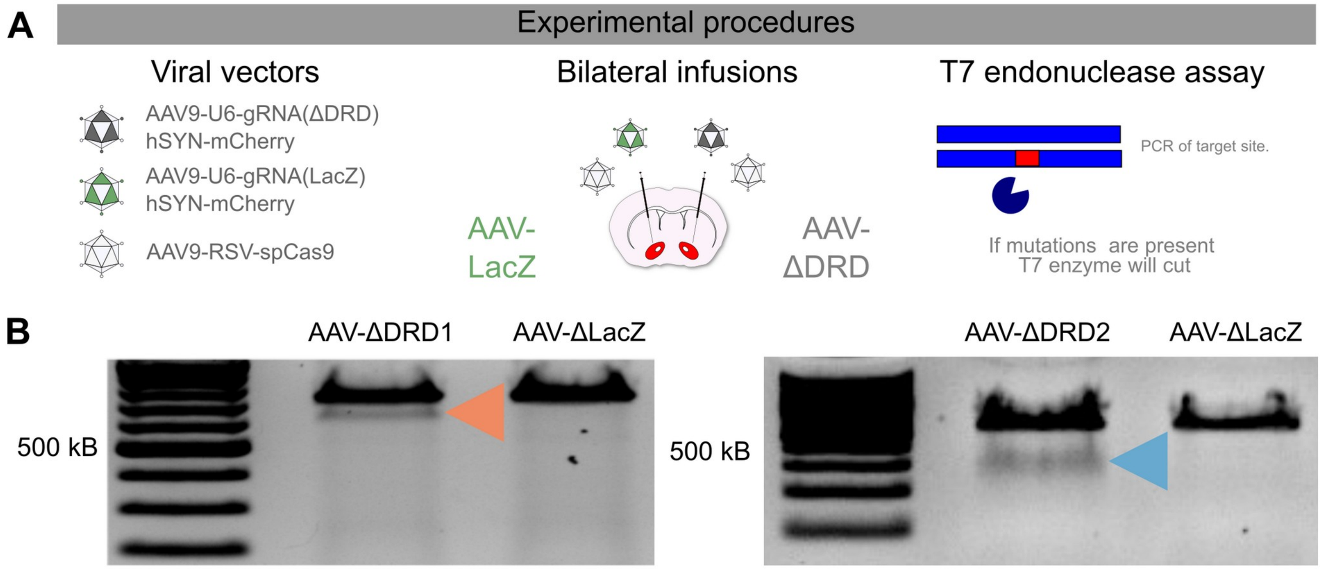


**Figure S1. Initial validation of AAV-CRISPR/Cas9 strategy to edit DA receptor genes in house mouse species. A)** Schematics of experimental procedure that the surgical approach and the assessment of gene editing. **B)** Images of agarose gel evidencing gene editing after infusion of AAV-ΔDRD1 and AAV-ΔDRD2, but not AAV-ΔLacZ

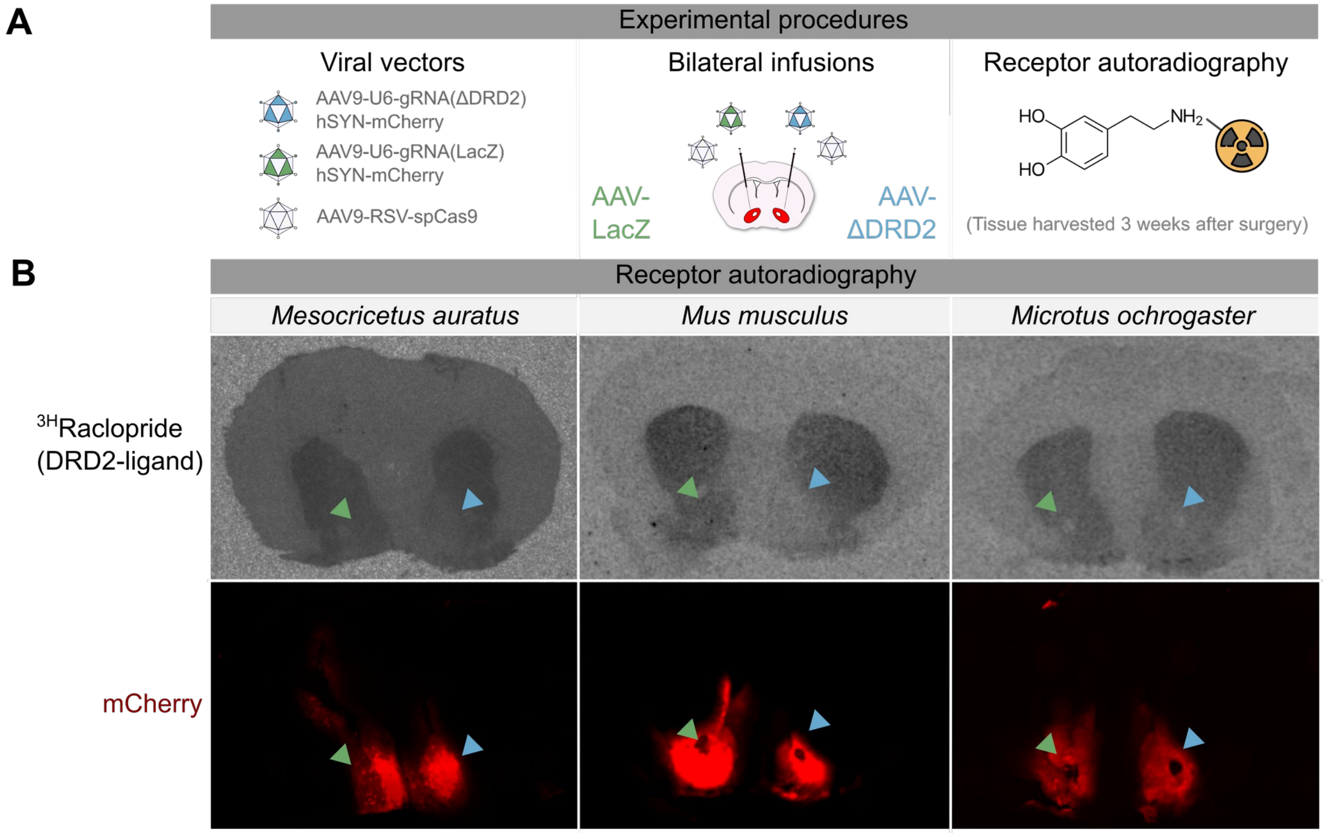


**Figure S2. Initial validation of AAV-CRISPR/Cas9 strategy to reduce DRD2 across rodent species.** (Top) Schematics of experimental procedure that the surgical approach and the validation of efficacy and selectivity of viral vectors by receptor autoradiography. (Bottom) Representative autoradiographies of 3HRaclopride binding, as well as images of the viral-induced mCherry expression counterstained. Green arrows indicate AAV-LacZ injections, while blue arrows represent AAV-ΔDRD2 infusions.
